# Temporal order of Alzheimer’s disease-related cognitive marker changes in BLSA and WRAP longitudinal studies

**DOI:** 10.1101/081174

**Authors:** Murat Bilgel, Rebecca L. Koscik, Yang An, Jerry L. Prince, Susan M. Resnick, Sterling C. Johnson, Bruno M. Jedynak

**Affiliations:** Laboratory of Behavioral Neuroscience, Intramural Research Program, National Institute on Aging,Baltimore, MD; CWisconsin Alzheimer’s Institute, University of Wisconsin School of Medicine and Public Health, USA, Madison, WI 53705; Department of Electrical and Computer Engineering, Johns Hopkins University, Baltimore, MD; Geriatric Research Education and Clinical Center, Wm. S. Middleton Veterans Hospital, USA, Madison WI 53705; Alzheimer’s Disease Research Center, University of Wisconsin School of Medicine and Public Health, USA, Madison, WI 53705; Waisman Laboratory for Brain Imaging and Behavior, University of Wisconsin-Madison, USA, Madison, WI 53705; Department of Mathematics and Statistics, Portland State University, Portland, OR, USA

**Author notes:** Corresponding author: Murat Bilgel, National Institute on Aging, Laboratory of Behavioral Neuroscience, 251 Bayview Blvd., Suite 100, Rm 04B329, Baltimore, MD 21224, USA.

**Keywords:** Digit span, progression score, Alzheimer’s disease, preclinical, BLSA, WRAP, cognitive measures

## Abstract

Investigation of the temporal trajectories of currently used neuropsychological tests is critical to identifying earliest changing measures on the path to dementia due to Alzheimer’s disease (AD). We used the Progression Score (PS) method to characterize the temporal trajectories of measures of verbal memory, executive function, attention, processing speed, language, and mental state using data spanning normal cognition, mild cognitive impairment (MCI), and AD from 1661 participants with a total of 7839 visits (age at last visit 77.6 SD 9.2) in the Baltimore Longitudinal Study of Aging and 1542 participants with a total of 4467 visits (age at last visit 59.9 SD 7.3) in the Wisconsin Registry for Alzheimer’s Prevention. This method aligns individuals in time based on the similarity of their longitudinal measurements to reveal temporal trajectories. As a validation of our methodology, we explored the associations between the individualized cognitive progression scores (Cog-PS) computed by our method and clinical diagnosis. Digit span tests were the first to show declines in both data sets, and were detected mainly among cognitively normal individuals. These were followed by tests of verbal memory, which were in turn followed by Trail Making Tests, Boston Naming Test, and Mini-Mental State Examination. Differences in Cog-PS across the clinical diagnosis groups were statistically significant, highlighting the potential use of Cog-PS as individualized indicators of disease progression. Identifying cognitive measures that are changing in preclinical AD can lead to the development of novel cognitive tests that are finely tuned to detecting earliest changes.

**ABBREVIATIONS:**

AD: Alzheimer’s disease
AVLT: Rey Auditory Verbal Learning Test
BLSA: Baltimore Longitudinal Study of Aging
CI: Cognitive impairment
Cog-PS: Cognitive progression score
CVLT: California Verbal Learning Test
AD: Alzheimer’s disease
MCI: Mild cognitive impairment
MMSE: Mini-Mental State Examination
WRAP: Wisconsin Registry for Alzheimer’s Prevention

## 1. INTRODUCTION

Cognitive changes in Alzheimer’s disease (AD), in particular declines in episodic memory, are detectable on neuropsychological testing up to fifteen years prior to clinical diagnosis [1–4]. However, cognition has only limited cross-sectional association with cerebral amyloid burden [5–11], which marks the beginning of preclinical AD according to the National Institute on Aging-Alzheimer’s Association (NIA–AA) criteria [12]. Longitudinal studies have consistently shown that amyloid levels are associated with greater rates of decline on tests of episodic memory [13–21], suggesting that amyloid changes precede episodic memory declines. These findings indicate that despite the fifteen-year period prior to diagnosis for detecting cognitive change, currently used neuropsychological tests fall short of detecting changes in the earliest disease stages where therapeutic intervention is hypothesized to be most promising. Therefore, there is a need to develop cognitive tests and batteries that are more sensitive to changes in early preclinical AD and that correlate better with AD-related neuroimaging measures. Cognitive measures that are dynamic in preclinical disease can facilitate clinical trials aimed at this early stage, as they can serve as outcome measures that are non-invasive and cheaper to administer than neuroimaging evaluations.

In order to develop such cognitive tests or batteries, it is necessary to study the neuroimaging correlates and temporal trajectories of currently used tests to identify earliest changing measures. Various studies have investigated correlations between cognitive measures and early AD-related neuroimaging markers [22–30]. In this work, we focus on characterizing the trajectories of a collection of cognitive markers widely used in studies of aging and AD. The ideal study of preclinical AD markers would follow cognitively normal individuals until they are diagnosed with dementia due to AD, and retrospectively analyze the time courses of the markers of those who developed AD. However, currently available sample sizes do not allow for cross-validation nor yield adequate statistical power to conduct such a study. To overcome this limitation, several statistical analysis approaches have been developed [31–36]. An important concept in a subset of these approaches is the time-alignment of individuals. Here, we used the Progression Score Model [36,37], a multivariate nonlinear mixed effects model, to construct cognitive marker trajectories spanning normal cognition, mild cognitive impairment (MCI), and AD stages by aligning individuals in time based on the similarity of their marker profiles. The method incorporates longitudinal information in performing this alignment and accounts for inter-individual differences in rate and baseline levels of progression. The main premise of the method is that rather than using age as a proxy for disease progression, we can obtain better disease progression indicators as well as temporal trajectories for the biomarkers by aligning individuals in time based on the similarity of their biomarker profiles. Time alignment of individuals allows us to study the long-term trajectories of cognitive measures despite the availability of only a small number of individuals who have traversed large extents of the cognitive trajectories. We conducted separate analyses on two well-characterized longitudinal studies, the Baltimore Longitudinal Study of Aging and the Wisconsin Registry for Alzheimer’s Prevention, in order to delineate the trajectories of measures of verbal memory, executive function, attention, processing speed, language, and mental state among individuals who are cognitively normal, have MCI, or have dementia due to AD.

The main aims of the analyses were to: 1) develop a standardized template of cognitive changes against which individuals can be quantified, 2) present evidence of the validity of the individualized scores obtained using the standardized cognitive template by exploring their associations with clinical diagnosis, 3) identify the order in which detectable changes begin to appear across cognitive tests, and 4) validate this identified ordering using an independent approach at the individual level.

## 2. METHODS

Statistical methods used in the following analyses were applied separately to two ongoing longitudinal cohort studies of human aging: the Baltimore Longitudinal Study of Aging (BLSA) [38], initiated in 1958 and conducted by the Intramural Research Program of the National Institute on Aging, and the Wisconsin Registry for Alzheimer’s Prevention (WRAP) [39], initiated in 2001 and conducted by the University of Wisconsin Alzheimer’s Institute.

### 2.1. Participants

#### BLSA participants

BLSA analyses were based on data from 1661 participants (Table 1). We included visits where participants were cognitively normal or had a clinical diagnosis of MCI or dementia due to AD. Baseline and last visit were defined for each participant as the first and last BLSA visit between 1/1992 and 11/2015 where they were aged ≥60 and met the inclusion criteria based on clinical diagnosis and number of available cognitive scores (see Section 2.2). A total of 7839 visits were selected for analysis. The Institutional Review Board of the NIA Intramural Research Program approved the research protocol for this study, and informed consent was obtained at each visit from all participants.

**Table 1.**
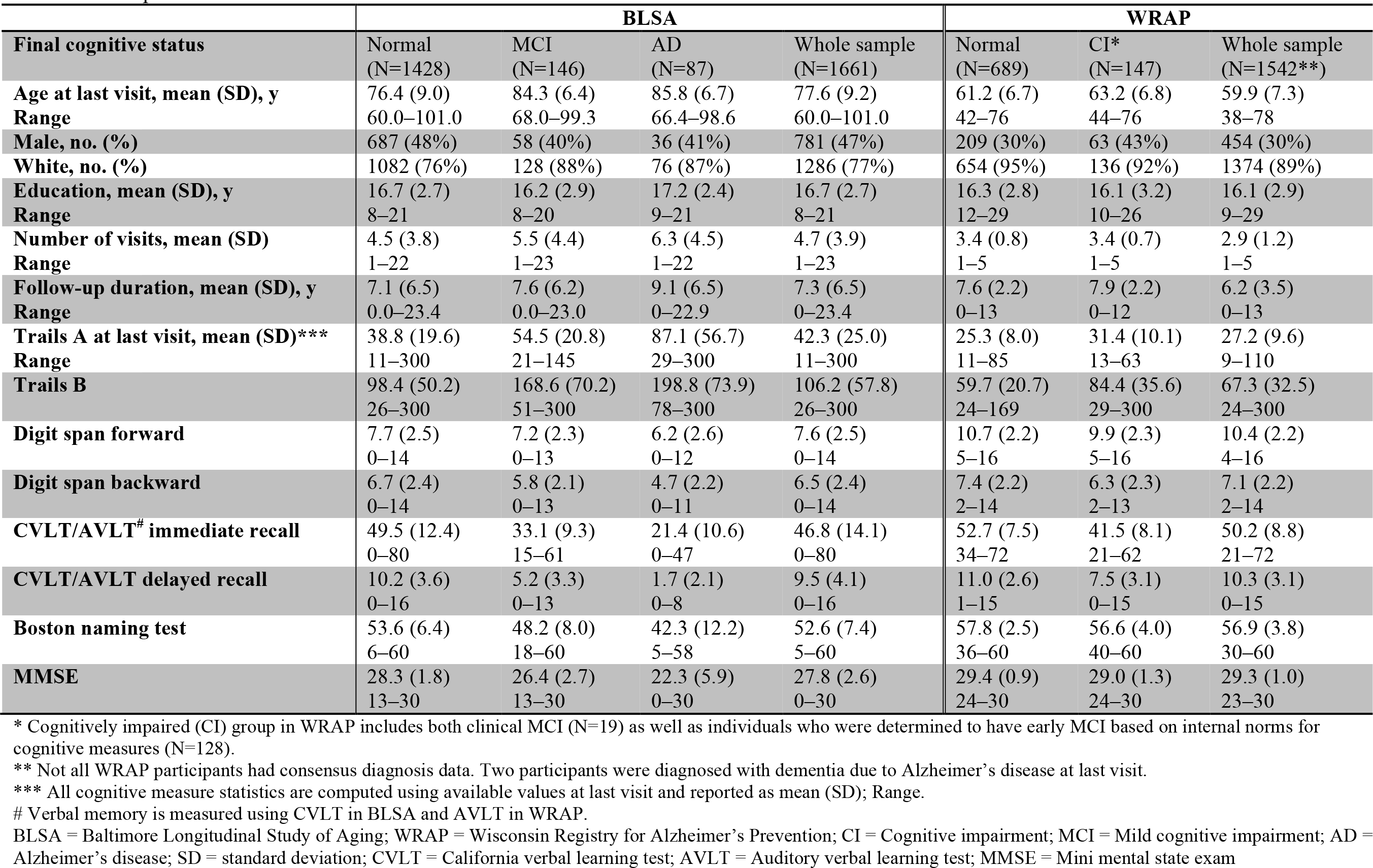
Participants.

|  | BLSA |  |  |  | WRAP |  |  |
| --- | --- | --- | --- | --- | --- | --- | --- |
| **Final cognitive status** | Normal<br>(N=1428) | MCI<br>(N=146) | AD<br>(N=87) | Whole sample<br>(N=1661) | Normal<br>(N=689) | CI*<br>(N=147) | Whole sample<br>(N=1542**) |
| **Age at last visit, mean (SD), y** | 76.4 (9.0) | 84.3 (6.4) | 85.8 (6.7) | 77.6 (9.2) | 61.2 (6.7) | 63.2 (6.8) | 59.9 (7.3) |
| **Range** | 60.0–101.0 | 68.0–99.3 | 66.4–98.6 | 60.0–101.0 | 42–76 | 44–76 | 38–78 |
| **Male, no. (%)** | 687 (48%) | 58 (40%) | 36 (41%) | 781 (47%) | 209 (30%) | 63 (43%) | 454 (30%) |
| **White, no. (%)** | 1082 (76%) | 128 (88%) | 76 (87%) | 1286 (77%) | 654 (95%) | 136 (92%) | 1374 (89%) |
| **Education, mean (SD), y** | 16.7 (2.7) | 16.2 (2.9) | 17.2 (2.4) | 16.7 (2.7) | 16.3 (2.8) | 16.1 (3.2) | 16.1 (2.9) |
| **Range** | 8–21 | 8–20 | 9–21 | 8–21 | 12–29 | 10–26 | 9–29 |
| **Number of visits, mean (SD)** | 4.5 (3.8) | 5.5 (4.4) | 6.3 (4.5) | 4.7 (3.9) | 3.4 (0.8) | 3.4 (0.7) | 2.9 (1.2) |
| **Range** | 1–22 | 1–23 | 1–22 | 1–23 | 1–5 | 1–5 | 1–5 |
| **Follow-up duration, mean (SD), y** | 7.1 (6.5) | 7.6 (6.2) | 9.1 (6.5) | 7.3 (6.5) | 7.6 (2.2) | 7.9 (2.2) | 6.2 (3.5) |
| **Range** | 0.0–23.4 | 0.0–23.0 | 0–22.9 | 0–23.4 | 0–13 | 0–12 | 0–13 |
| **Trails A at last visit, mean (SD)\*\*\*** | 38.8 (19.6) | 54.5 (20.8) | 87.1 (56.7) | 42.3 (25.0) | 25.3 (8.0) | 31.4 (10.1) | 27.2 (9.6) |
| **Range** | 11–300 | 21–145 | 29–300 | 11–300 | 11–85 | 13–63 | 9–110 |
| **Trails B** | 98.4 (50.2) | 168.6 (70.2) | 198.8 (73.9) | 106.2 (57.8) | 59.7 (20.7) | 84.4 (35.6) | 67.3 (32.5) |
|  | 26–300 | 51–300 | 78–300 | 26–300 | 24–169 | 29–300 | 24–300 |
| **Digit span forward** | 7.7 (2.5) | 7.2 (2.3) | 6.2 (2.6) | 7.6 (2.5) | 10.7 (2.2) | 9.9 (2.3) | 10.4 (2.2) |
|  | 0–14 | 0–13 | 0–12 | 0–14 | 5–16 | 5–16 | 4–16 |
| **Digit span backward** | 6.7 (2.4) | 5.8 (2.1) | 4.7 (2.2) | 6.5 (2.4) | 7.4 (2.2) | 6.3 (2.3) | 7.1 (2.2) |
|  | 0–14 | 0–13 | 0–11 | 0–14 | 2–14 | 2–13 | 2–14 |
| **CVLT/AVLT<sup>#</sup> immediate recall** | 49.5 (12.4) | 33.1 (9.3) | 21.4 (10.6) | 46.8 (14.1) | 52.7 (7.5) | 41.5 (8.1) | 50.2 (8.8) |
|  | 0–80 | 15–61 | 0–47 | 0–80 | 34–72 | 21–62 | 21–72 |
| **CVLT/AVLT delayed recall** | 10.2 (3.6) | 5.2 (3.3) | 1.7 (2.1) | 9.5 (4.1) | 11.0 (2.6) | 7.5 (3.1) | 10.3 (3.1) |
|  | 0–16 | 0–13 | 0–8 | 0–16 | 1–15 | 0–15 | 0–15 |
| **Boston naming test** | 53.6 (6.4) | 48.2 (8.0) | 42.3 (12.2) | 52.6 (7.4) | 57.8 (2.5) | 56.6 (4.0) | 56.9 (3.8) |
|  | 6–60 | 18–60 | 5–58 | 5–60 | 36–60 | 40–60 | 30–60 |
| **MMSE** | 28.3 (1.8) | 26.4 (2.7) | 22.3 (5.9) | 27.8 (2.6) | 29.4 (0.9) | 29.0 (1.3) | 29.3 (1.0) |
|  | 13–30 | 13–30 | 0–30 | 0–30 | 24–30 | 24–30 | 23–30 |

\* Cognitively impaired (CI) group in WRAP includes both clinical MCI (N=19) as well as individuals who were determined to have early MCI based on internal norms for cognitive measures (N=128).
\*\* Not all WRAP participants had consensus diagnosis data. Two participants were diagnosed with dementia due to Alzheimer's disease at last visit.
\*\*\* All cognitive measure statistics are computed using available values at last visit and reported as mean (SD); Range.

#### WRAP participants

WRAP analyses were based on data from 1542 participants (Table 1), 72.3% of whom had a family history of AD. We included all visits where participants were cognitively normal or had a clinical diagnosis of MCI (including an “early MCI” diagnosis, defined in Section 2.3) or dementia due to AD. Baseline and last visit were defined for each participant as the first and last WRAP visit up until Wave 5 (inclusive) where they met the inclusion criteria based on clinical diagnosis and number of available cognitive scores (see Section 2.2). A total of 4467 visits were selected for analysis. All activities for this study were approved by the Institutional Review Board and completed in accordance with the Helsinki Declaration.

### 2.2. Cognitive measures

Eight cognitive measures were selected for our analyses: Trail Making Tests (Trails) A and B [40] to assess processing speed and executive function, Wechsler Adult Intelligence Scale (WAIS) digit span forward and backward (revised edition [41] in BLSA, 3^rd^ edition [42] in WRAP) to assess attention and executive function, California Verbal Learning Test (CVLT) [43] in BLSA and Rey Auditory Verbal Learning Test (AVLT) [44] in WRAP measuring immediate recall (sum of total recall across five learning trials) and 20-minute delayed recall to assess verbal memory, Boston Naming Test [45] to assess language, and Mini-Mental State Examination (MMSE) [46] to assess mental state. Trails A and B were truncated at 300 seconds. All measures were kept in their original scales. We included visits where at least four of these eight measures were available. We did not include WAIS digit symbol substitution [41,42] or animal fluency [47] in our analyses even though they were common to both data sets, since they were introduced in later stages in BLSA and/or WRAP.

CVLT was added to the BLSA battery in 3/1993 and digit span was added in 1/1992, while the remainder of the measures were added between 5/1984 and 1/1990. BLSA participants under age 60 do not receive the Boston naming test as part of their cognitive battery and did not receive the Trails making test until 2005; therefore, these test scores were not included. MMSE was added to the WRAP battery in Wave 2, while the remaining eight measures were available starting with Wave 1 in 2001.

### 2.3. Clinical Diagnoses

#### BLSA methods

All BLSA participants are reviewed for cognitive impairment at a consensus case conference if they have a score ≥4 on the Blessed Information–Memory– Concentration Test [48], if their Clinical Dementia Rating (CDR) score [49] is ≥0.5 on the subject or informant report or if they screen abnormal on the Dementia Questionnaire. Consensus diagnoses are determined by a research team that includes neurologists, psychiatrists, neuropsychologists, research nurses, and research assistants. Diagnoses of dementia and AD are based on Diagnostic and Statistical Manual of Mental Disorders 3^rd^ edition–revised [50] and National Institute of Neurological and Communication Disorders and Stroke–AD and Related Disorders Association (NINCDS-ADRDA) [51] criteria, respectively. Mild cognitive impairment (MCI) is diagnosed based on the Petersen criteria [52] when cognitive impairment is evident for a single domain or multiple domains without significant functional loss in activities of daily living. Out of the eight cognitive measures described in Section 2.2, only Trails A and B, Boston Naming, and MMSE are used in clinical diagnosis determination.

#### WRAP methods

WRAP participant visits are reviewed at a consensus case conference if they meet one or more of the following criteria: 1) cognitive abnormalities relative to WRAP peers (i.e., at least 1.5 standard deviations below expected relative to robust internal norms adjusting for age, sex, and literacy-level on the most recent assessment for factor scores or individual measures of memory, executive function, language, working memory, or attention [53], [54]; 2) cognitive performance on one or more tests below values used in other studies as cutpoints for mild cognitive impairment (MCI) diagnoses (e.g., WMS-R Logical Memory II [55] story A score <9, Alzheimer’s Disease Neuroimaging Initiative [56]); or 3) an abnormal informant report indicating subjective cognitive or functional decline. Consensus diagnoses are determined by a research team that includes physicians, clinical neuropsychologists, and clinical nurse practitioners.

Cognitive statuses include cognitively normal, early MCI, clinical MCI, other cognitive impairment (e.g., due to other medical conditions), or dementia. The diagnosis of clinical MCI is based on NIA-AA criteria [57] and includes a) concern regarding change in cognition, b) impairment in one or more cognitive domains, c) preservation of functional abilities, and d) does not meet criteria for dementia. The status of early MCI was developed to identify individuals in the cohort who exhibit lower than expected objective performance in one or more cognitive domains (relative to internal robust norms), but may not yet report subjective cognitive complaints. This experimental construct is thought to represent a phenotype of early cognitive decline expected to precede a clinical diagnosis of MCI [54,58]. Dementia diagnosis is based on NINCDS-ADRDA criteria [51]. Both demented WRAP individuals included in these analyses had dementia due to AD. The consensus review process was initiated in late 2012. At the time of these analyses, consensus diagnoses were available for each participant’s last attended study visit.

### 2.4. Statistical analyses

#### 2.4.1. Progression score model

We used an improved version [37] of the Progression Score (PS) method [36] to compute cognitive progression scores (Cog-PS) for individuals in BLSA and WRAP in separate models using the eight cognitive tests described in Section 2.2 (see Supplementary Material for details). The Progression Score method is based on a multivariate model that enables the computation of a score for each visit using a collection of longitudinal biomarker measures to reflect the state of the visit relative to the rest of the sample. Cog-PS is an affine transformation of age, and these transformations are allowed to vary across individuals via subject-specific variables that model inter-subject differences in rate and baseline levels of progression. The scale of the Cog-PS at the end of the model fitting procedure is arbitrary (i.e., a Cog-PS of 0 does not convey any meaning on its own, but compared to a Cog-PS of 1, it indicates better overall performance on the cognitive measures included in the model). In order to give meaning to the Cog-PS scale, we shifted all Cog-PS values such that a Cog-PS of 0 corresponds to the mean value across cognitively normal individuals, and then scaled all Cog-PS values such that the standard deviation of Cog-PS among cognitively normal individuals is 1. The Cog-PS calibration, when combined with the appropriate transformations of the model parameters, does not affect the likelihood of the model (i.e., the calibrated and uncalibrated models are identical in terms of their fit to the observed data). This calibration is intended to make Cog-PS values computed on separate data sets comparable, with the underlying assumption that cognitively normal groups are comparable. At the end of model fitting, we obtained Cog-PS for each visit as well as estimates of cognitive measure trajectories as a function of Cog-PS. The overall procedure is summarized in Figure S1.

#### 2.4.2. Temporal ordering of cognitive measures

To compare the estimated trajectories for the eight cognitive measures, we linearly scaled them such that their estimated values at the minimum and maximum Cog-PS observed in the sample were 0 and 1, respectively. In this standardized space, we refer to a scaled cognitive value of 0 as “normal” and a scaled cognitive value of 1 as “abnormal”. We visually evaluated the estimated trajectories to make determinations about temporal ordering of cognitive changes. To complement this qualitative evaluation, we developed a procedure that relies on a threshold for each cognitive measure to obtain quantifiable measures to determine temporal ordering. We transform the midway point of 0.5 in this standardized space back into the unstandardized scale for each cognitive measure to obtain a threshold for each cognitive measure. These cognitive measure thresholds are not intended for classification of individuals. They rather serve as tools for understanding the order in which changes begin to appear across cognitive measures. To this end, we determined the Cog-PS value at which the estimated trajectories surpass the cognitive measure thresholds. Cognitive measures whose trajectories surpass the threshold at a lower Cog-PS value are measures that change earlier in the disease process. In order to provide further evidence for the ordering of cognitive marker changes, we performed 20 bootstrap experiments. The number of bootstrap experiments was limited to 20 due to the time consuming model fitting procedure. We visualized the resulting trajectories and compared the Cog-PS values at which measures crossed the threshold across the bootstrap experiments to quantify the statistical confidence associated with this ordering.

#### 2.4.3. Validation of Cog-PS results

To show evidence for the validity of the individualized Cog-PS, we first investigated differences in Cog-PS cross-sectionally across diagnosis groups using two-sided Wilcoxon rank-sum tests. We used the last visit for this analysis since diagnosis was available only at last visit in WRAP.

To provide further evidence for the temporal ordering of cognitive marker changes, we used an analysis based on a multi-state Markov model that was independent from the Cog-PS approach. In this analysis, we used the mean values of the cognitive markers within the MCI or CI ([M]CI) group at last visit (listed in Table 1) as thresholds for categorizing each cognitive measure at each visit as being normal or abnormal, with the exception of MMSE, where we defined abnormality using the conservative threshold ≤ 25 rather than using the sample mean since there is a strong ceiling effect in our samples. For Trails A and B, abnormality was defined as being equal to or exceeding the threshold, and for all other cognitive measures as being equal to or below the threshold. We considered each pair of cognitive measures, denoted *x* and *y*, in separate Markov models. We categorized each visit into one of the following four states: *x* and *y* both normal (state 1), only *x* abnormal (state 2A), only *y* abnormal (state 2B), *x* and *y* both abnormal (state 3). If both *x* and *y* were missing, the visit was excluded from analysis. If only one was available, then the state of the visit was censored accordingly (i.e., if only *x* is available and is normal, then the state is either 1 or 2B). We assumed that individuals need to pass through one of the intermediate states 2A or 2B in order to transition between states 1 and 3. All other transitions, including backward transitions, were included in the model, which is summarized in Figure S2. We fitted this multi-state Markov model using the msm [59] package in R (version 3.2.1) [60]. We are interested in comparing the transitions out of the first state where both measures are normal. To this end, we compared the transition rates into States 2A and 2B from State 1.

## 3. RESULTS

### 3.1. Temporal ordering of cognitive measures

Estimated cognitive marker trajectories as function of Cog-PS are presented in Figure 1. While digit span tests have a marked downward slope in the left-hand side of the figure as a Cog-PS of –2, other cognitive measures do not begin exhibiting marked declines until later along the Cog-PS scale. CVLT/AVLT measures are next to exhibit declines, followed by Trails, Boston Naming, and MMSE.

**Figure 1.**
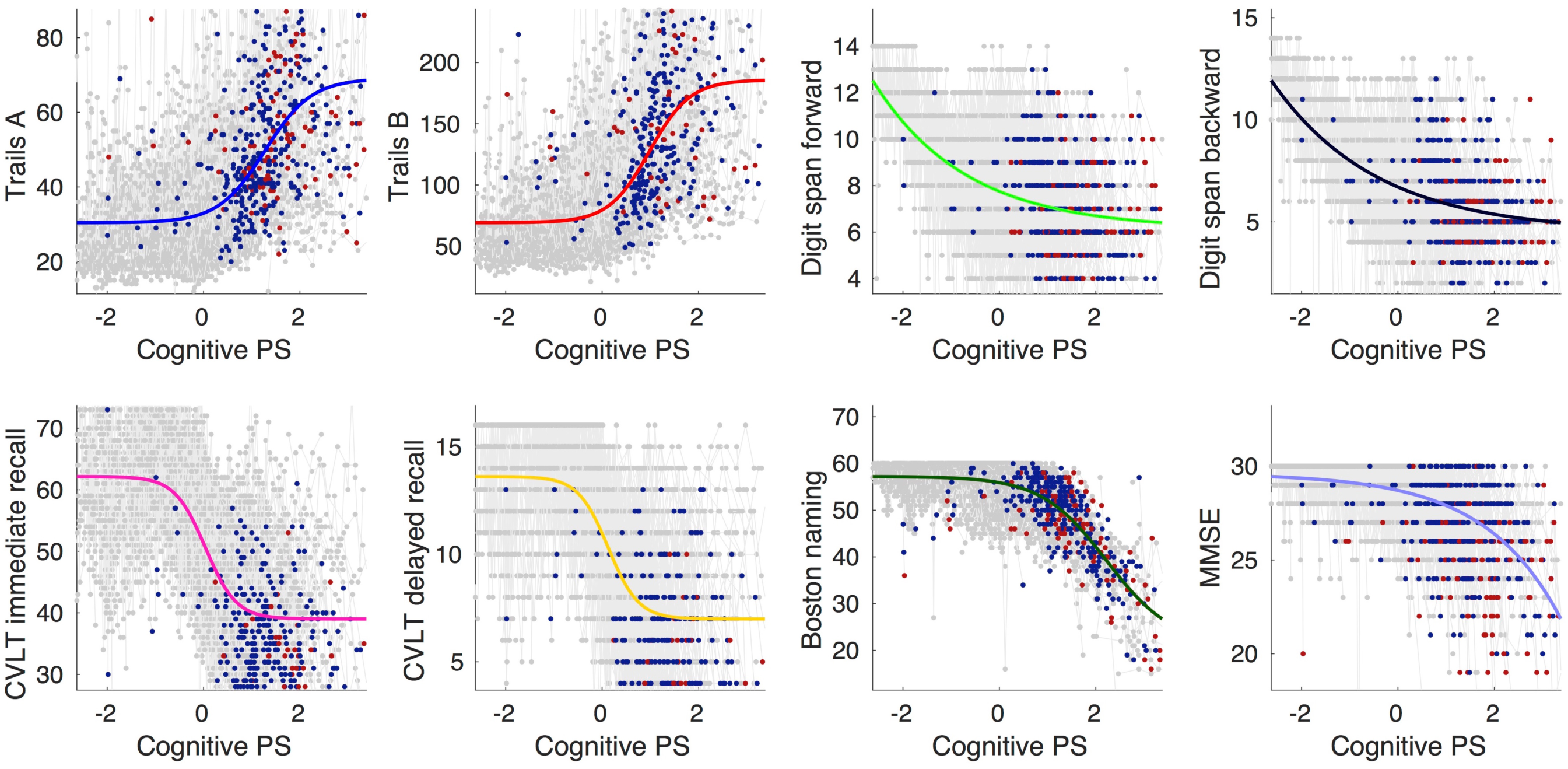
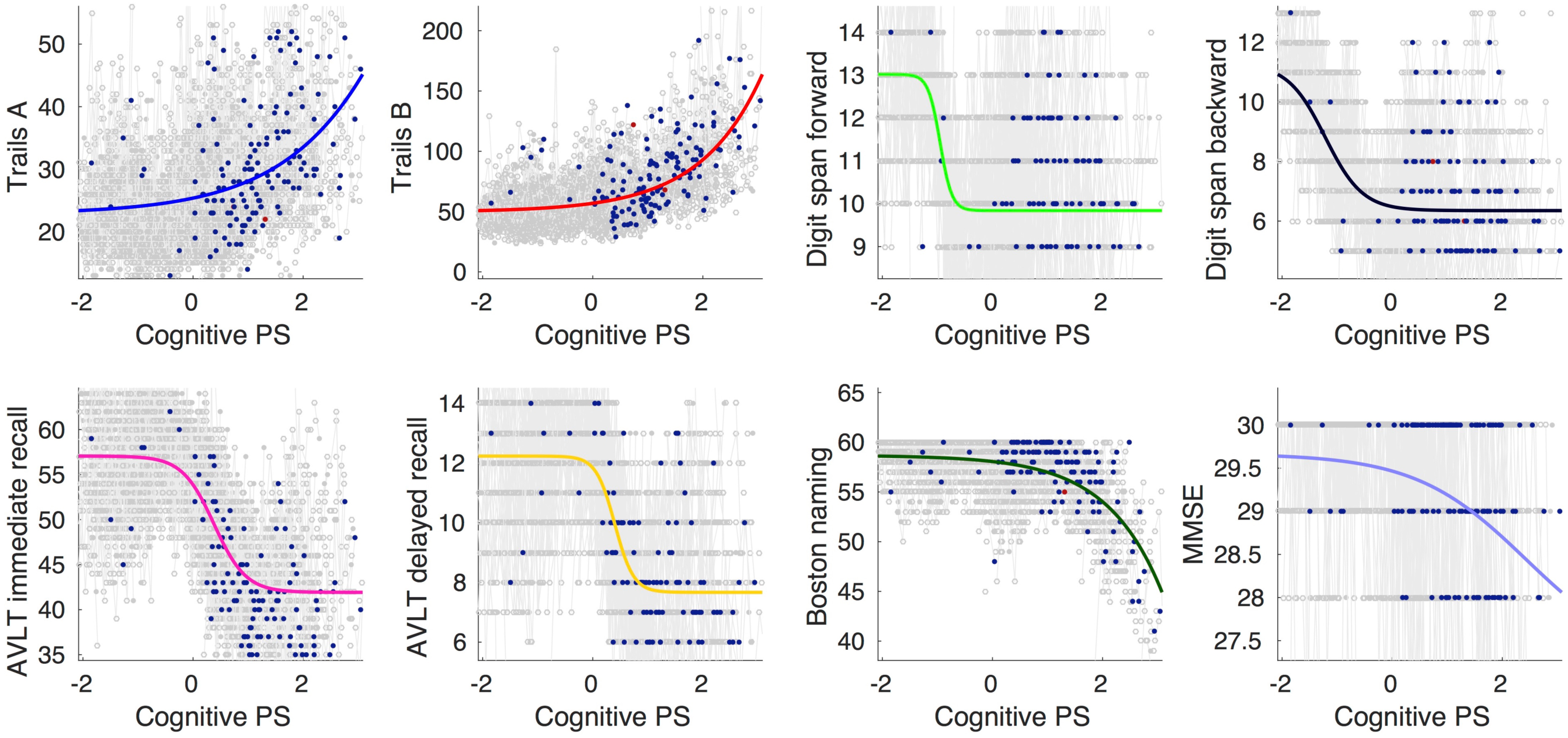
Cognitive measure trajectories in (a) BLSA and (b) WRAP. Gray, blue, and red dots indicate cognitively normal, MCI or CI ([M]CI), and AD visits, respectively. Visits where consensus diagnoses are not available are indicated as unfilled circles.

The standardized cognitive template, obtained by scaling the cognitive measures based on fitted values corresponding to the minimum and maximum Cog-PS values in the sample, is presented in Figure 2. The cognitive thresholds obtained by transforming the midway point along the *y*-axis back into the unstandardized scale for each cognitive marker are presented in Table 2, along with the Cog-PS values at which the estimated trajectories attain these thresholds. In both BLSA and WRAP, digit span forward and backward were the first measures to attain the cognitive thresholds, followed by CVLT/AVLT immediate and delayed recall, and finally Trails A and B, Boston naming, and MMSE.

**Figure 2.**
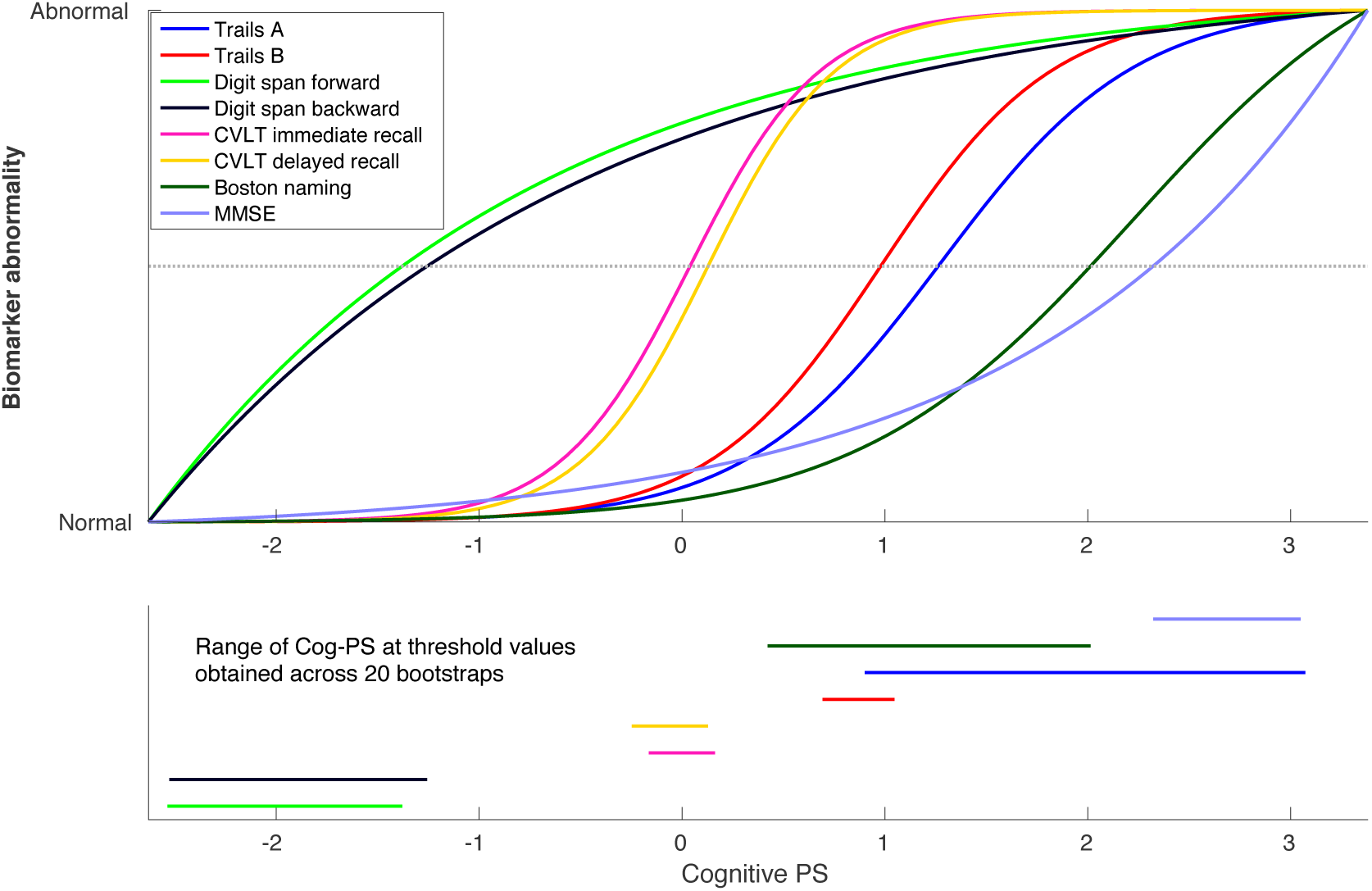
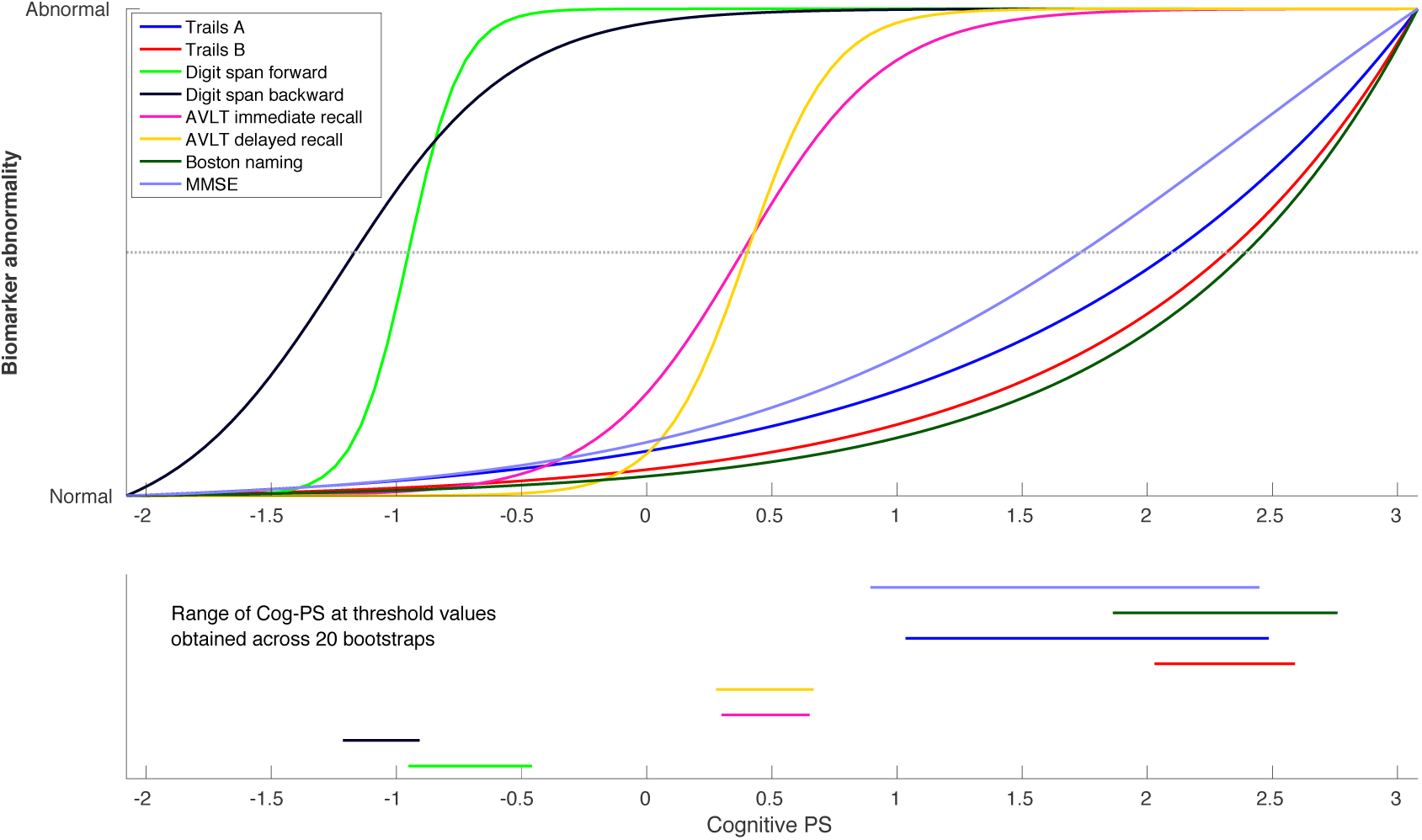
Scaled cognitive trajectories obtained using (a) BLSA and (b) WRAP data. Dotted line in the top panel corresponds to the midway point between Normal and Abnormal. Line segments in the bottom panel indicate the range of the Cog-PS values that attain the midway point (i.e., cognitive threshold) across 20 bootstrap experiments.

**Table 2.**
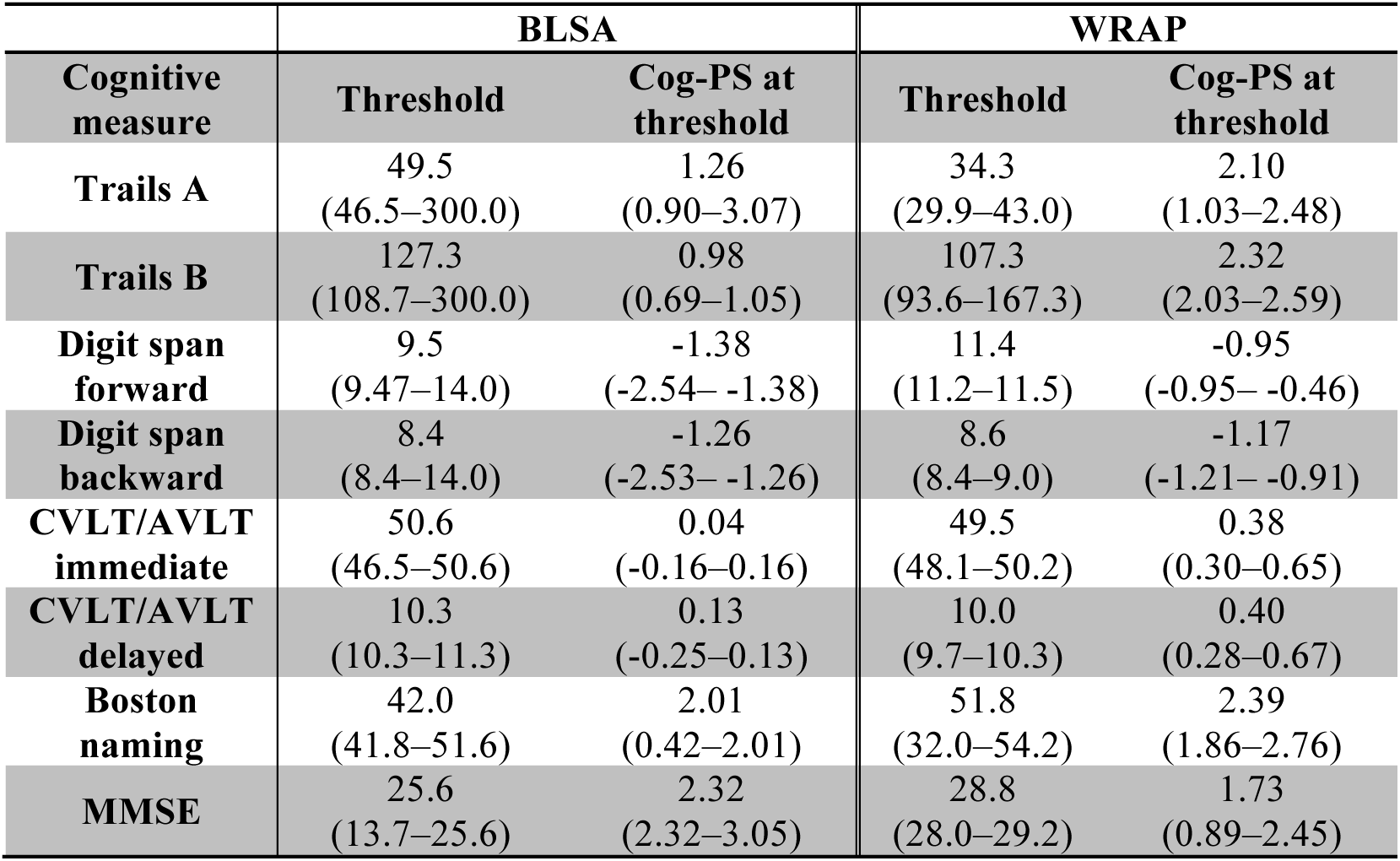
Cognitive measure thresholds computed on the whole sample. Minimum and maximum values obtained across 20 bootstraps are indicated in parentheses.

| Cognitive measure | BLSA |  | WRAP |  |
| --- | --- | --- | --- | --- |
|  | Threshold | Cog-PS at threshold | Threshold | Cog-PS at threshold |
| **Trails A** | 49.5<br>(46.5–300.0) | 1.26<br>(0.90–3.07) | 34.3<br>(29.9–43.0) | 2.10<br>(1.03–2.48) |
| **Trails B** | 127.3<br>(108.7–300.0) | 0.98<br>(0.69–1.05) | 107.3<br>(93.6–167.3) | 2.32<br>(2.03–2.59) |
| **Digit span forward** | 9.5<br>(9.47–14.0) | -1.38<br>(-2.54– -1.38) | 11.4<br>(11.2–11.5) | -0.95<br>(-0.95– -0.46) |
| **Digit span backward** | 8.4<br>(8.4–14.0) | -1.26<br>(-2.53– -1.26) | 8.6<br>(8.4–9.0) | -1.17<br>(-1.21– -0.91) |
| **CVLT/AVLT immediate** | 50.6<br>(46.5–50.6) | 0.04<br>(-0.16–0.16) | 49.5<br>(48.1–50.2) | 0.38<br>(0.30–0.65) |
| **CVLT/AVLT delayed** | 10.3<br>(10.3–11.3) | 0.13<br>(-0.25–0.13) | 10.0<br>(9.7–10.3) | 0.40<br>(0.28–0.67) |
| **Boston naming** | 42.0<br>(41.8–51.6) | 2.01<br>(0.42–2.01) | 51.8<br>(32.0–54.2) | 2.39<br>(1.86–2.76) |
| **MMSE** | 25.6<br>(13.7–25.6) | 2.32<br>(2.32–3.05) | 28.8<br>(28.0–29.2) | 1.73<br>(0.89–2.45) |

### 3.2. Validation of Cog-PS results

Cog-PS at last visit was associated with concurrent diagnosis, with cognitively normal individuals having lower Cog-PS compared to impaired individuals (Figure 3). All pairwise group comparisons (normal vs. [M]CI, MCI vs. AD, and normal vs. AD) were significant within each data set (two-sided Wilcoxon rank-sum test, all *p*<.00002).

**Figure 3.**
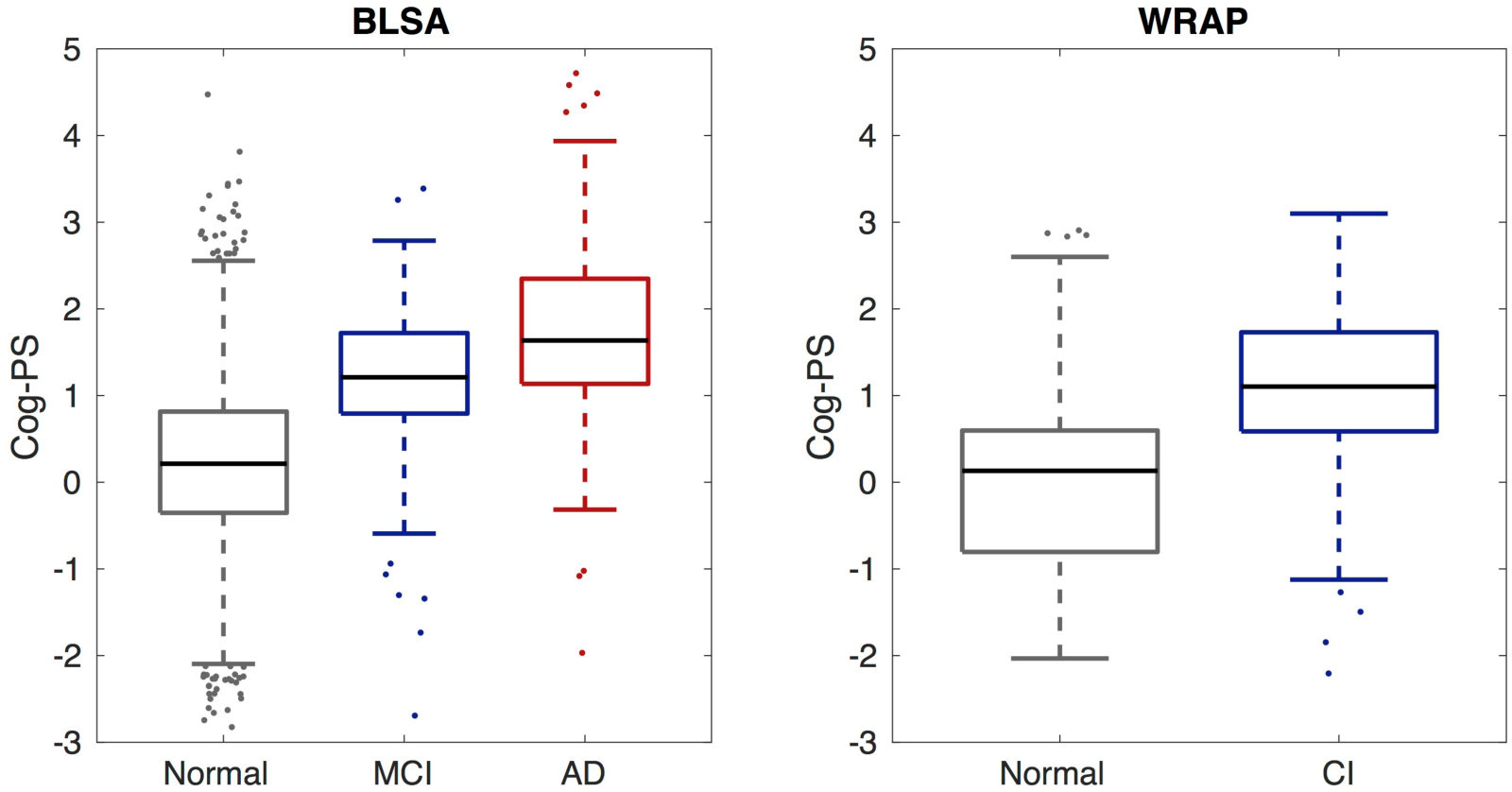
Box plot of Cog-PS by clinical diagnosis at last visit in BLSA (left) and WRAP (right). Central mark is the median, edges of the box correspond to the interquartile range, and whiskers extend to the range of non-outlier Cog-PS values. Outliers are plotted individually. All pairwise group comparisons were significant within each data set (two-sided Wilcoxon rank-sum test, *p* < 0.00002).

Estimated transition rates using the Markov model (Table S2) suggest that cognitive measures attain the mean level observed in the [M]CI group in the following order in BLSA: Digit span backward, digit span forward, CVLT immediate, CVLT delayed, Trails A, Trails B, MMSE, and Boston naming. In WRAP, the ordering is as follows: digit span forward, digit span backward, AVLT delayed, AVLT immediate, Trails A, Trails B, and MMSE. Results in WRAP suggest that change in Boston naming occurs after digit span backward and before MMSE, but its exact placement among AVLT and Trails is not clear.

## 4. DISCUSSION

We applied the progression score model to align individuals in time based on the similarity of their cognitive marker profiles to learn long-term cognitive trajectories from shorter-term longitudinal measurements. From this model, we obtained individualized scores, termed Cog-PS, which we validated by demonstrating their association with clinical diagnosis. Our analyses were conducted separately on data from two well-characterized longitudinal studies of aging, BLSA and WRAP, and revealed similar results.

Using the estimated trajectories, we showed that digit span tests exhibit their most extensive dynamic range while individuals are cognitively normal, and that these changes precede those observed on verbal memory tests, which in turn precede changes on Trails, Boston naming, and MMSE. The trajectory estimates we obtained using the progression score model along with their approximate confidence intervals as illustrated in Figure 2 demonstrated this temporal ordering of the cognitive measures. We also used an approach based on a threshold for each cognitive measure to quantify and complement the findings of the qualitative assessment.

Our findings about ordering of cognitive changes were validated using a multistate Markov model. While the Markov model confirmed our finding that digit span measures are the first to exhibit changes, followed by changes in CVLT/AVLT, its results regarding temporal ordering individual cognitive measures were not as consistent between the two data sets as the Cog-PS model. These discrepancies may be due to the sensitivity of the Markov model to the cognitive thresholds used to define the states in the model.

The progression score approach enabled the computation of an individualized indicator of cognitive performance based on a collection of longitudinal measurements. We demonstrated that there are statistically significant differences in progression score across cognitively normal, [M]CI, and AD groups. Since the progression score model is agnostic to clinical diagnosis, this analysis served as a validation of the computed scores.

However, it should be noted that this is a partial validation since there is a degree of circularity due to the fact that four out of the eight cognitive measures were used in diagnosis determination in BLSA, and eight were used in WRAP.

While we used a calibration approach to render the progression scores computed on BLSA and WRAP comparable, it is important to note that due to differences in the composition of cognitively normal groups in the two data sets, Cog-PS values do not convey the same meaning across the studies. Despite this limitation of our methodology, we found similar temporal cognitive patterns in both studies. Cognitively normal as well as the [M]CI groups in BLSA had lower cognitive performance on all tests compared to WRAP. Our scaling procedure for establishing correspondences across cognitive markers is sensitive to the range of cognitive measurements present in the study. Since this scaling procedure is intended to identify normal and abnormal cognitive test values, it is important that the data set contain measurements spanning the entire normal-to-abnormal range for each cognitive test. This condition is not fully satisfied in either data set we considered in this work, particularly in WRAP since WRAP participants are about 18 years younger than BLSA participants on average. Therefore, the scaling procedure may be less accurate in WRAP than in BLSA, especially for later-changing measures such as Boston naming and MMSE.

Several previous studies have reported that changes in digit span tests are detectable prior to a clinical diagnosis of cognitive impairment. For example, a study that evaluated non-demented individuals with subjective memory complaints found that those with normal digit span scores (as defined using age- and education-adjusted neuropsychological test scores) at baseline did not exhibit significant declines on verbal memory, visual memory, or executive function after a mean follow-up of 6.6 years, but had significant declines on the sum score of digit span forward and backward [61]. On the other hand, age-, sex-, and education-matched individuals with impaired digit span scores at baseline had significant declines on tests of verbal learning and animal fluency but not on any other cognitive test. These findings are in agreement with our estimated ordering of cognitive trajectories on the path to cognitive impairment, with digit span measures declining first, followed by measures of verbal memory. Another study that investigated changes in cognition prior to autopsy in a sample of individuals who remained cognitively normal found that while longitudinal changes were not significant when assessed separately among individuals with and without AD neuropathology, longitudinal decline in the attention/working memory domain (assessed using digit span forward and backward tests) was greater among those with neuropathology compared to those without [62]. Other domains, including episodic memory, language, and executive function, did not show statistically significant longitudinal differences between the two groups [62].

A limitation of our method is that it does not indicate whether the detected cognitive changes are due to AD-related mechanisms, and further studies are needed to delineate normal aging versus disease. Therefore, our results cannot be interpreted as evidence for using digit span to predict individualized diagnoses. Digit span is not a good predictor of concurrent diagnosis; using cognitive measure thresholds based on our Cog-PS model to classify individuals as normal versus [M]CI/AD yields a large number of false positives. This lack of specificity of digit span for dementia has been documented previously [63,64]. What our results demonstrate is that changes in digit span are most evident early on in cognitively normal stages. Detectability of these changes and their associations with future outcomes at the individual-level, including AD diagnosis in following years or decades, remain to be fully elucidated; however, prior studies have reported that sensitivity to change on digit span is small [65]. Another limitation of our method is that it treats discrete cognitive measures such as digit span as continuous variables, and therefore may introduce bias into the characterization of their longitudinal trajectories.

Despite these limitations, understanding the association of digit span tests with brain changes can be informative for designing novel cognitive tests or batteries that are more sensitive to changes in preclinical AD and that correlate with functional and structural brain changes over the course of disease.

Several studies of a small number of healthy individuals found that higher performance on digit span backward is associated with activation of the right dorsolateral prefrontal cortex (DLPFC) [66–68], bilateral inferior parietal lobule, anterior cingulate, left DLPFC, and Broca’s area, with a subset of these regions also implicated in relation to digit span forward [67,68]. DLPFC is one of the amyloid accumulating regions, and therefore it may be possible to demonstrate associations between digit span performance and DLPFC amyloid levels among cognitively normal individuals in studies with large sample sizes.

Analyses conducted in parallel in multiple data sets or in combined samples will accelerate efforts to further elucidate the relationships among cognitive measures implicated in preclinical AD. Identifying cognitive measures that are dynamic in preclinical AD can lead to the development of novel cognitive tests that are finely tuned to detecting earliest changes. Such measures will facilitate clinical trials aimed at this early stage by serving as outcome measures that are non-invasive and cheaper to administer than neuroimaging.

## ACKNOWLEDGMENTS

We would like to thank the BLSA and WRAP participants and staff for their efforts and commitment. Without them this research would not have been possible.

This research was supported in part by the Intramural Research Program of the National Institute on Aging (National Institutes of Health) and the Michael J. Fox Foundation for Parkinson’s Research, MJFF Research Grant ID: 9310 (BMJ). WRAP is supported by NIA grant R01AG27161 (SCJ; Wisconsin Registry for Alzheimer Prevention: Biomarkers of Preclinical AD). WRAP is also supported by the Clinical and Translational Science Award (CTSA) program, through the NIH National Center for Advancing Translational Sciences (NCATS), grant UL1TR000427. The content is solely the responsibility of the authors and does not necessarily represent the official views of the NIH.

## SUPPLEMENTARY MATERIAL

### Progression Score Model

For clarity, vector-valued variables are in bold and matrices are capitalized. The affine transformation between the age *t*_*ij*_ of subject *i* at visit *j* and the Cog-PS *s*_*ij*_ is given by

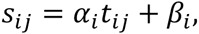

where (*α*_*i*_ and *β*_*i*_ are the subject-specific variables assumed to be independent and identically distributed across subjects with a bivariate normal distribution *N*(**m**, *V*). *α*_*i*_ and *β*_*i*_ model the rate of change and baseline level of Cog-PS, respectively.

The trajectory of cognitive measure *k* is assumed to be a sigmoid in Cog-PS, and is given by

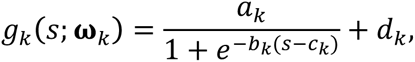

where **ω**_*k*_ = (*a*_*k*_, *b*_*k*_, *c*_*k*_,*d*_*k*_) are trajectory parameters to be estimated. *d*_*k*_ and *a*_*k*_ + *d*_*k*_ correspond to the minimum and maximum values of the sigmoid, respectively. *c*_*k*_ is the inflection point (the Cog-PS value at which the second derivative is zero) and *a*_*k*_*b*_*k*_/4 is the slope at the inflection point.

The observed cognitive measures *y*_*ijk*_ stacked into the vector **y**_*ij*_ are assumed to have additive normally distributed noise, and are described by

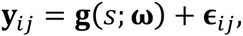

where **g** is the vector obtained by stacking *g*_*k*_, and **ϵ**_*ij*_ is noise, assumed to be independent and identically distributed with a multivariate normal distribution *N*(**0**, *R*). *R* is an unstructured covariance matrix that represents the variance of noise for each cognitive measure as well as the correlations among them, and is estimated during the model fitting procedure.

Model fitting is performed using a Monte Carlo expectation-maximization (MC-EM) algorithm. The subject-specific variables and missing cognitive measures constitute the hidden variables in this framework. Model parameters include **ω**, **m**, *V*, and *R*. The EM approach is an iterative procedure where the most likely values of the hidden variables are computed given the data and current parameter estimates, and then the model parameters are estimated using these most likely values for the hidden variables. Since the integral in the E-step for our model does not have an analytical form, we approximate it using Monte Carlo samples.

After model fitting, we compute the cross-sectional mean and variance of the Cog-PS among cognitively normal individuals:

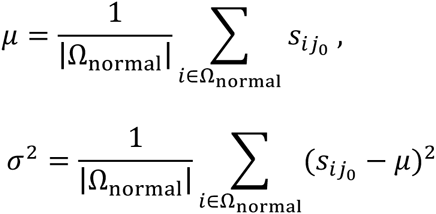

where *j*_*0*_ is the visit index at which the mean and variance are computed and Ω_normal_ is the set of individuals who are cognitively normal at visit *j*_*0*_. For BLSA, *j*_*0*_ corresponds to the baseline visit, and for WRAP to the last visit (since diagnosis information is available only at last visit). We calibrate the Cog-PS scale as 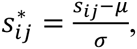 which corresponds to the following changes in the subject-specific variables:

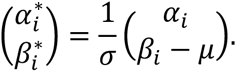

This calibration is accompanied by the following standardization of the model parameters:

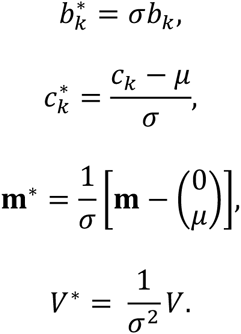

Let the minimum and maximum progression scores observed in the data set after model fitting be *s*_min_ and *s*_max_. We scale the trajectory of each marker so that fitted values at these values correspond across markers. Scaled values are given by

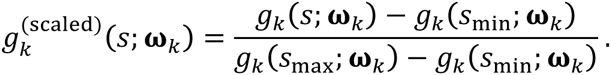

**Figure S1.**
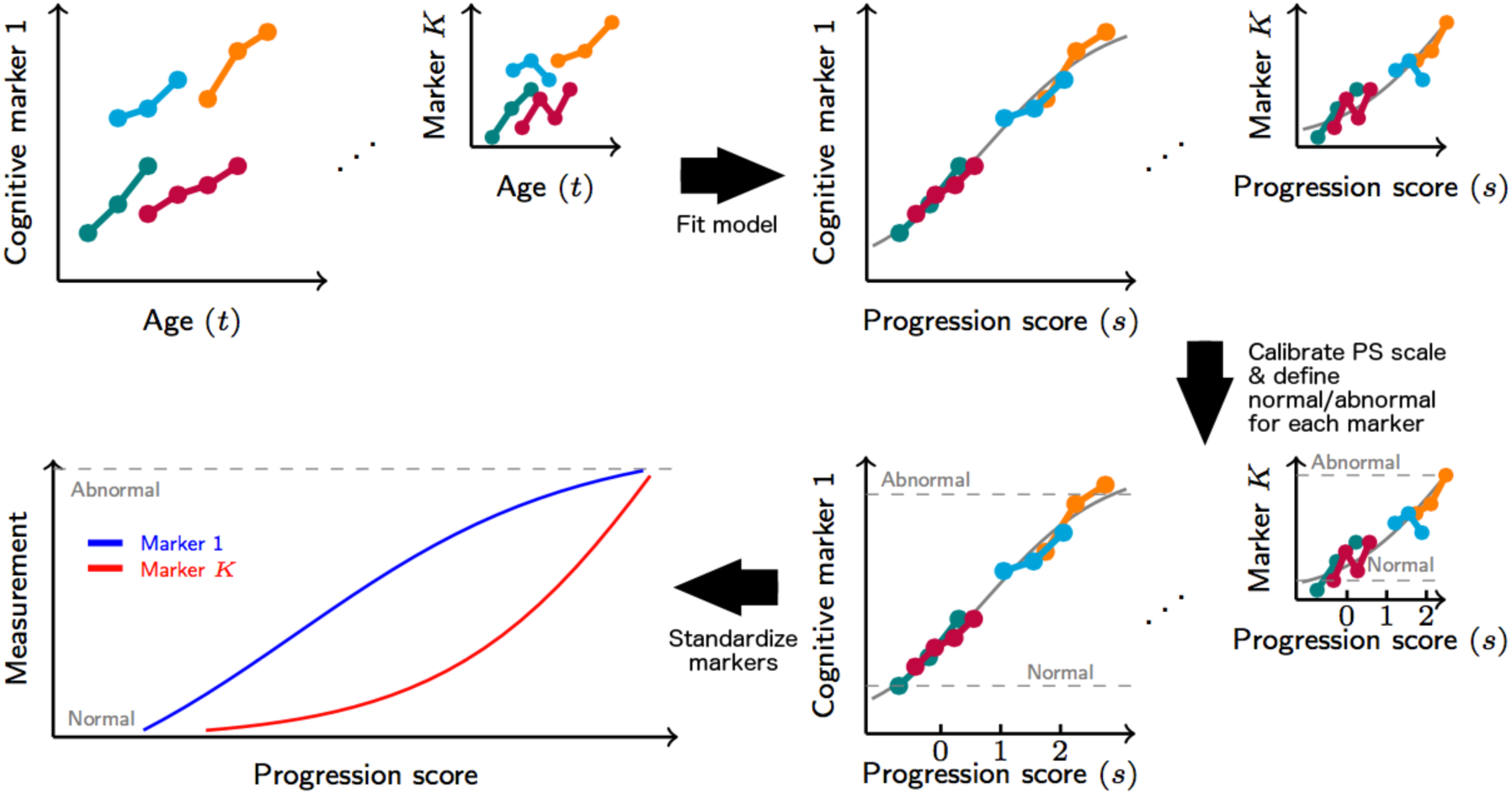
Diagram illustrating progression score (PS) model fitting, PS calibration, and cognitive measurement scaling to obtain standardized space of cognitive markers. Lower values for the illustrated cognitive markers indicate lower cognitive performance. PS values are calibrated such that lower progression scores indicate better overall cognitive performance.

### Multi-state Markov model

**Figure S2.**
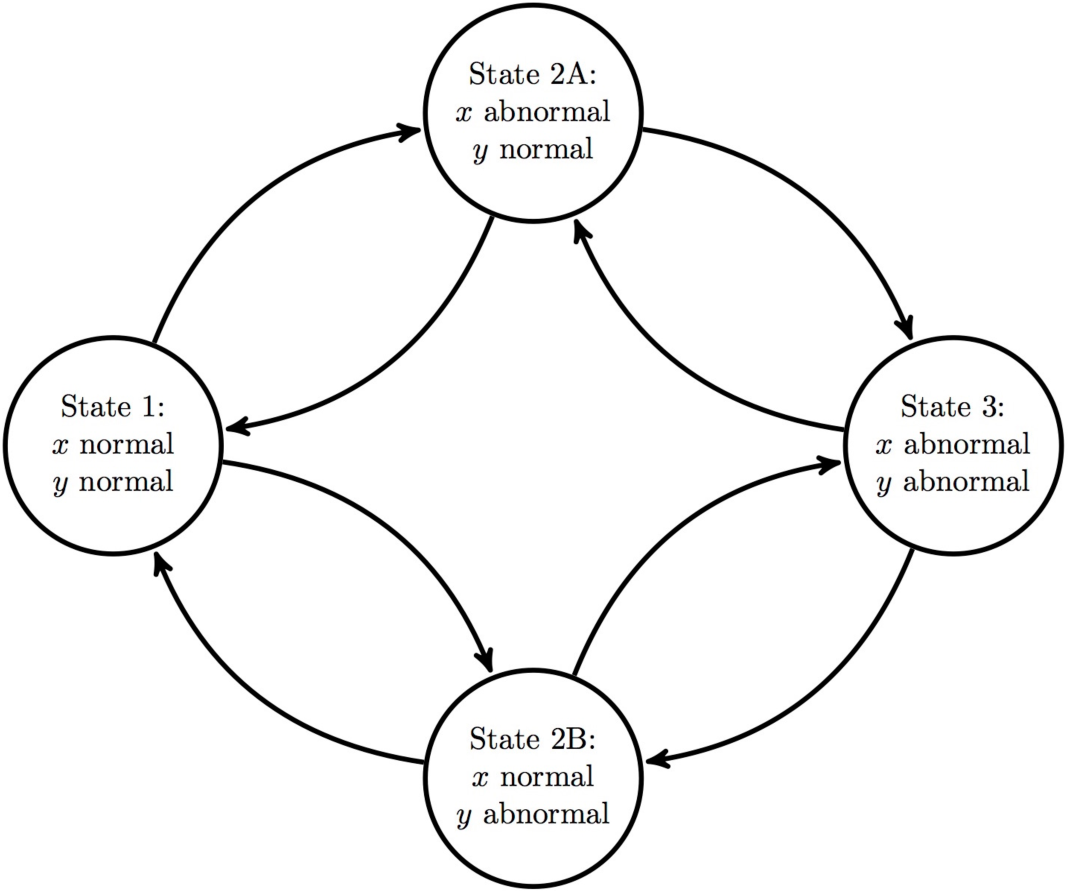
Multi-state Markov model. Each node corresponds to one of the four states in the model. *x* and *y* correspond to cognitive measures. Normal and abnormal categorization of the cognitive measures are based on thresholding at the mean value within the [M]CI group at last visit.

We present detailed results comparing digit span forward and CVLT delayed recall tests in BLSA. Table S1 presents the counts of consecutive states in the BLSA, and Figure S3 displays the estimated transition rates. Since we did not characterize transition rates by age, our results represent the transition rates for an individual at the sample mean age.

**Table S1.**
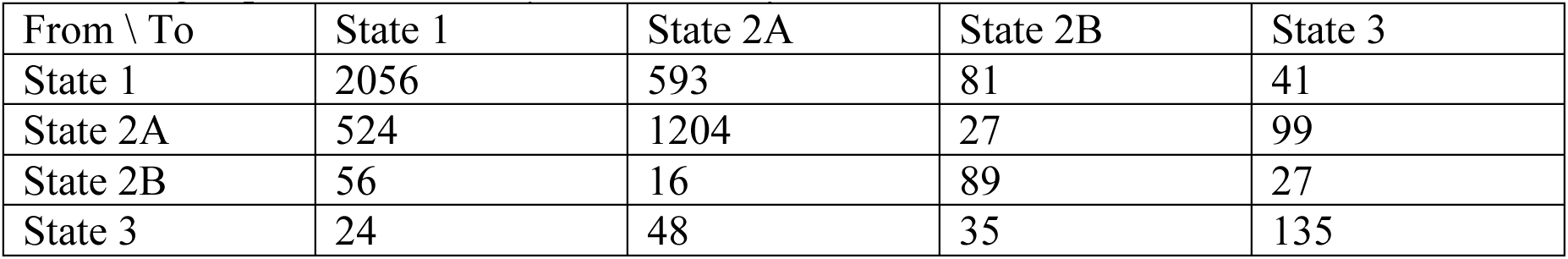
Summary of the BLSA data as a frequency table of pairs of consecutive states for *x*=Digit span forward and *y*=CVLT delayed recall.

| From \ To | State 1 | State 2A | State 2B | State 3 |
| --- | --- | --- | --- | --- |
| State 1 | 2056 | 593 | 81 | 41 |
| State 2A | 524 | 1204 | 27 | 99 |
| State 2B | 56 | 16 | 89 | 27 |
| State 3 | 24 | 48 | 35 | 135 |

**Figure S3.**
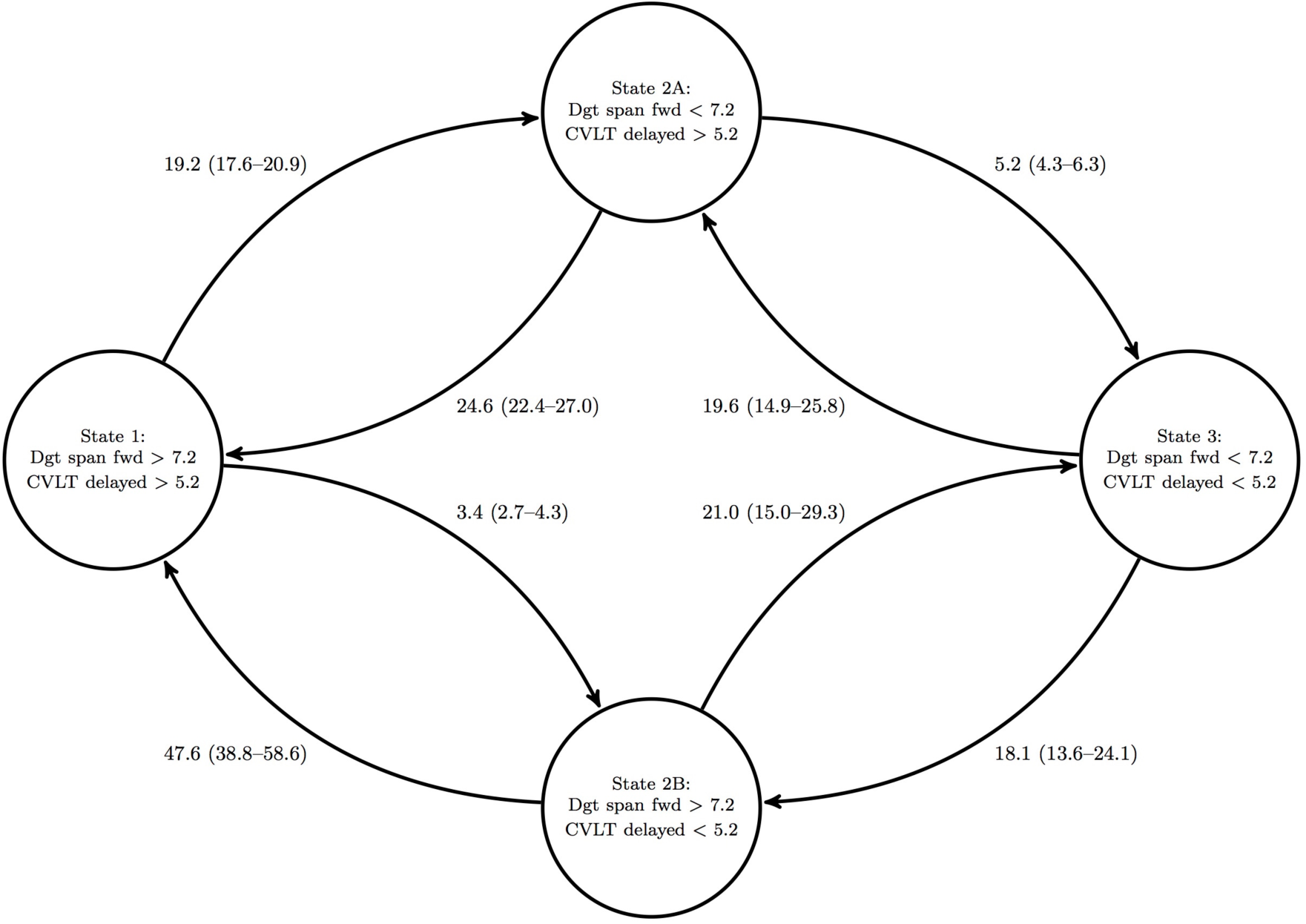
Multi-state Markov model for *x*=Digit span forward and *y*=CVLT delayed recall. Transition rates per 100 person-years, estimated using BLSA data, are shown next to the arrows. 95% confidence intervals for the transition rates are in parentheses.

Assuming an initial cohort consisting of 100 individuals in State 1 at baseline, the model estimates that approximately 53 of them would be in State 1, 37 in State 2A, 4 in State 2B, and 6 in State 3 after a follow-up of 7.3 years (the mean follow-up duration in the BLSA).

Transition rates out of state 1 are shown in Table S2 for all Markov models fitted using each pair (*x,y*) of cognitive tests.

**Table S2.** State transition rates per 100 person-years. 95% confidence intervals are in parentheses. The higher transition rate is in bold whenever it is statistically different from the lower rate based on 95% CIs. If the State 1 → 2A transition rate is higher, the measure listed under column *x* becomes abnormal before *y*. If the State 1 → 2B transition rate is higher, the measure listed under column *y* becomes abnormal before *x*.

| Cognitive measure pair (x,y) |  | BLSA |  | WRAP |  |
| --- | --- | --- | --- | --- | --- |
| x | y | State 1 → 2A | State 1 → 2B | State 1 → 2A | State 1 → 2B |
| Trails A | Trails B | **4.20**<br>(3.57–4.95) | 3.28<br>(2.70–3.98) | **6.79**<br>(5.77–8.00) | 2.88<br>(2.22–3.75) |
|  | Digit span forward | 4.62<br>(3.74–5.72) | **18.89**<br>(17.31–20.62) | 6.65<br>(5.54–7.98) | 7.23<br>(6.23–8.40) |
|  | Digit span backward | 4.33<br>(3.63–5.16) | **11.21**<br>(10.10–12.43) | 6.11<br>(4.95–7.54) | **11.47**<br>(9.94–13.24) |
|  | CVLT/AVLT immediate | 4.38<br>(3.76–5.10) | 4.10<br>(3.50–4.80) | **6.75**<br>(5.75–7.92) | 2.78<br>(2.23–3.48) |
|  | CVLT/AVLT delayed | 4.25<br>(3.66–4.93) | 3.74<br>(3.20–4.38) | **7.04**<br>(5.99–8.26) | 3.39<br>(2.79–4.13) |
|  | Boston naming | **4.82**<br>(4.15–5.60) | 2.46<br>(2.08–2.91) | **7.19**<br>(6.07–8.50) | 4.16<br>(3.49–4.96) |
|  | MMSE | **4.95**<br>(4.32–5.67) | 2.70<br>(2.19–3.32) | **7.40**<br>(6.46–8.48) | 0.42<br>(0.15–1.18) |
| Trails B | Digit span forward | 2.94<br>(2.21–3.91) | **19.21**<br>(17.63–20.93) | 3.18<br>(2.48–4.07) | **7.19**<br>(6.27–8.24) |
|  | Digit span backward | 3.00<br>(2.41–3.73) | **10.89**<br>(9.83–12.07) | 3.52<br>(2.65–4.68) | **11.81**<br>(10.39–13.43) |
|  | CVLT/AVLT immediate | 4.04<br>(3.41–4.78) | 3.72<br>(3.15–4.39) | 3.17<br>(2.58–3.89) | 3.12<br>(2.59–3.76) |
|  | CVLT/AVLT delayed | **3.99**<br>(3.37–4.73) | 3.24<br>(2.74–3.84) | 3.63<br>(2.99–4.41) | 3.40<br>(2.83–4.08) |
|  | Boston naming | **4.17**<br>(3.54–4.91) | 2.30<br>(1.94–2.74) | 4.32<br>(3.51–5.31) | 3.60<br>(3.00–4.32) |
|  | MMSE | **4.08**<br>(3.49–4.76) | 2.28<br>(1.84–2.83) | **4.48**<br>(3.83–5.24) | 0.2<br>(0.06–0.75) |
| Digit span forward | Digit span backward | **16.98**<br>(15.30–18.83) | 8.10<br>(6.67–9.83) | 4.79<br>(3.81–6.02) | **9.52**<br>(8.05–11.26) |
|  | CVLT/AVLT immediate | **18.77**<br>(17.21–20.47) | 3.43<br>(2.68–4.41) | **7.08**<br>(6.18–8.11) | 3.31<br>(2.67–4.11) |
|  | CVLT/AVLT delayed | **19.21**<br>(17.63–20.93) | 3.42<br>(2.71–4.31) | **7.16**<br>(6.21–8.25) | 3.68<br>(3.02–4.49) |
|  | Boston naming | **18.64**<br>(17.07–20.36) | 2.06<br>(1.60–2.67) | **7.31**<br>(6.32–8.46) | 3.64<br>(2.94–4.50) |
|  | MMSE | **19.13**<br>(17.58–20.81) | 2.81<br>(1.93–4.11) | **7.72**<br>(6.83–8.73) | 0.45<br>(0.12–1.74) |
| Digit span backward | CVLT/AVLT immediate | **11.55**<br>(10.43–12.78) | 3.86<br>(3.14–4.75) | **11.21**<br>(9.82–12.79) | 3.14<br>(2.42–4.06) |
|  | CVLT/AVLT delayed | **11.75**<br>(10.62–13.01) | 3.48<br>(2.86–4.23) | **11.04**<br>(9.66–12.62) | 3.62<br>(2.88–4.54) |
|  | Boston naming | **11.53**<br>(10.39–12.80) | 2.12<br>(1.72–2.61) | **11.48**<br>(9.99–13.19) | 3.40<br>(2.68–4.30) |
|  | MMSE | **11.82**<br>(10.70–13.06) | 3.52<br>(2.66–4.65) | **12.06**<br>(10.73–13.57) | 0.48<br>(0.17–1.33) |
| CVLT/AVLT immediate | CVLT/AVLT delayed | 3.35<br>(2.58–4.37) | 2.78<br>(2.17–3.57) | 2.82<br>(2.16–3.69) | 2.90<br>(2.29–3.67) |
|  | Boston naming | **3.58**<br>(3.07–4.19) | 2.14<br>(1.79–2.55) | 3.17<br>(2.59–3.87) | **3.81**<br>(3.21–4.52) |
|  | MMSE | **4.38**<br>(3.75–5.12) | 3.13<br>(2.54–3.86) | **3.72**<br>(3.19–4.33) | 0.07<br>(0.01–0.56) |
| CVLT/AVLT delayed | Boston naming | **3.77**<br>(3.24–4.38) | 2.12<br>(1.78–2.53) | 3.40<br>(2.80–4.13) | 3.85<br>(3.24–4.59) |
|  | MMSE | **3.87**<br>(3.31–4.52) | 2.92<br>(2.35–3.63) | **4.06**<br>(3.50–4.71) | 0.16<br>(0.04–0.71) |
| Boston naming | MMSE | 2.44<br>(2.07–2.86) | 2.92<br>(2.36–3.60) | **4.43**<br>(3.82–5.14) | 0.41<br>(0.15–1.12) |

